# A DNA part library for reliable engineering of the emerging model nematode symbiotic bacterium *Xenorhabdus griffiniae* HGB2511

**DOI:** 10.1101/2025.06.09.658710

**Authors:** Elin M. Larsson, Olivia Y. Wang, Richard M. Murray

## Abstract

*Xenorhabdus griffiniae* is a bacterium that lives inside the intestine of the entomopathogenic nematode *Steinernema hermaphroditum* and partners with the nematode to infect and kill insect larvae in soil. The construction of gene circuits, like reporters, in *X. griffiniae* would provide tools to study and better understand the symbiotic relationship it has with its host. However, because *X. griffiniae* is not a model organism, information about gene circuit construction in *X. griffiniae* is limited. We develop and characterize a DNA part library similar to the CIDAR MoClo extension library for *E. coli* to allow more efficient construction of genetic circuits in *X. griffiniae*. TurboRFP expressing strains with different constitutive Anderson promoters and different ribosome binding sites (RBS) were constructed to quantify promoter and RBS strengths in *X. griffiniae*. Furthermore, two fluorescent proteins sfGFP and sfYFP, as well as the bioluminescent *luxCDABE* operon were added to the part library and successfully expressed in *X. griffiniae*. We then used the characterized parts to build and characterize IPTG inducible constructs.

## Introduction

Nematodes are gaining increasing interest as simple models for studying gut microbiome interactions within a host [1–4]. The entomopathogenic nematodes of genus *Steinernema* are emerging as a model system because they often exhibit highly species-specific associations with a single bacterial strain of *Xenorhabdus* spp. to constitute its core microbiome. The emerging nematode model *Steinernema hermaphroditum* and its symbiotic bacterium *X. griffiniae* are of particular interest due to the nematode being consistently hermaphroditic and both partners being genetically tractable [5–8]. The bacterial symbionts are important for the development of the nematodes by providing nutrients [9] as well as a crucial part of their insect-killing life style. The bacteria make a cocktail of insect immunosuppressants, toxins and antibiotics to aid in killing the larvae and protecting the cadaver from the surrounding soil microbiome [10]. Their ability to kill insect larvae together has shed light on them as potential bioalternatives supplementing chemical pesticides in combating crop pests in agriculture.

There are many open questions about how this mutualistic relationship emerges and is maintained. Many of these knowledge gaps persist because of a lack of a genetic toolkit to develop synthetic genetic circuits such as biosensors and actuators within *Xenorhabdus*, to enable sensing and modulation of the nematode gut environment. Well-characterized libraries of genetic parts with different gene expression properties are an essential component of such genetic toolkits [11, 12].

Unlike model organisms like *E. coli*, the number of available well-characterized DNA parts for engineering *X. griffiniae* is small. To our knowledge, there are only two fluorescent proteins, one ribosome binding site (RBS), and one constitutive promoter described in *X. griffiniae* [8]. Here, we build and characterize a genetic part library consisting of constitutive promoters, RBS, and coding sequences for *X. griffiniae*. The *in vitro* characterization of the parts translates to relative performance in the *in vivo* context and allow predictable construction of IPTG inducible constructs in *X. griffiniae* for the first time.

The donor strains have been submitted to Addgene and can be used to integrate these constructs in other *Xenorhabdus* species or to make the vector needed to construct new genetic circuits using the parts characterized in this work.

## Results

### Characterization of DNA libraries in *Xenorhabdus griffiniae*

To enable rapid cloning followed by conjugation into *Xenorhabdus*, we modified an existing Tn7-based conjugation plasmid (HGB1262)[8] by adding UNS sequences flanking the insert region of the plasmid. By doing this, the plasmid backbone can be used for 3G cloning [13] with the CIDAR MoClo kit [14] and the CIDAR MoClo extension kit [15]. Using this plasmid, genetic constructs can be integrated at a single locus on the *X. griffiniae* chromosome, thereby avoiding variability in plasmid copy number.

We choose to focus on promoters and RBS because they allow tuning of the expression of fluorescent proteins that are already known to express well in *X. griffiniae*. We built promoter and RBS constructs by varying only the promoter or RBS and fixing the RBS or promoter respectively (Figure 1A). After building the promoter and RBS libraries, we characterized their relative strengths by measuring the fluorescence output normalized by optical density (OD) after 24 hours of growth. The strongest promoter is P5d (J23102), which is consistent with the strength in *E. coli* [14] (Figure 1B), and the strongest RBS is U25m (Figure 1C). The U25m RBS sequence is a variant of U4m that includes the genetic insulator RiboJ upstream. Previous work in *E. coli* has shown that RiboJ increases the expression of downstream genes by between 2-fold and 10-fold [16]. Here, we see that RiboJ increases the expression of TurboRFP driven by U4m by nearly 7-fold, which is consistent with the *E. coli* data.

**Figure 1.**
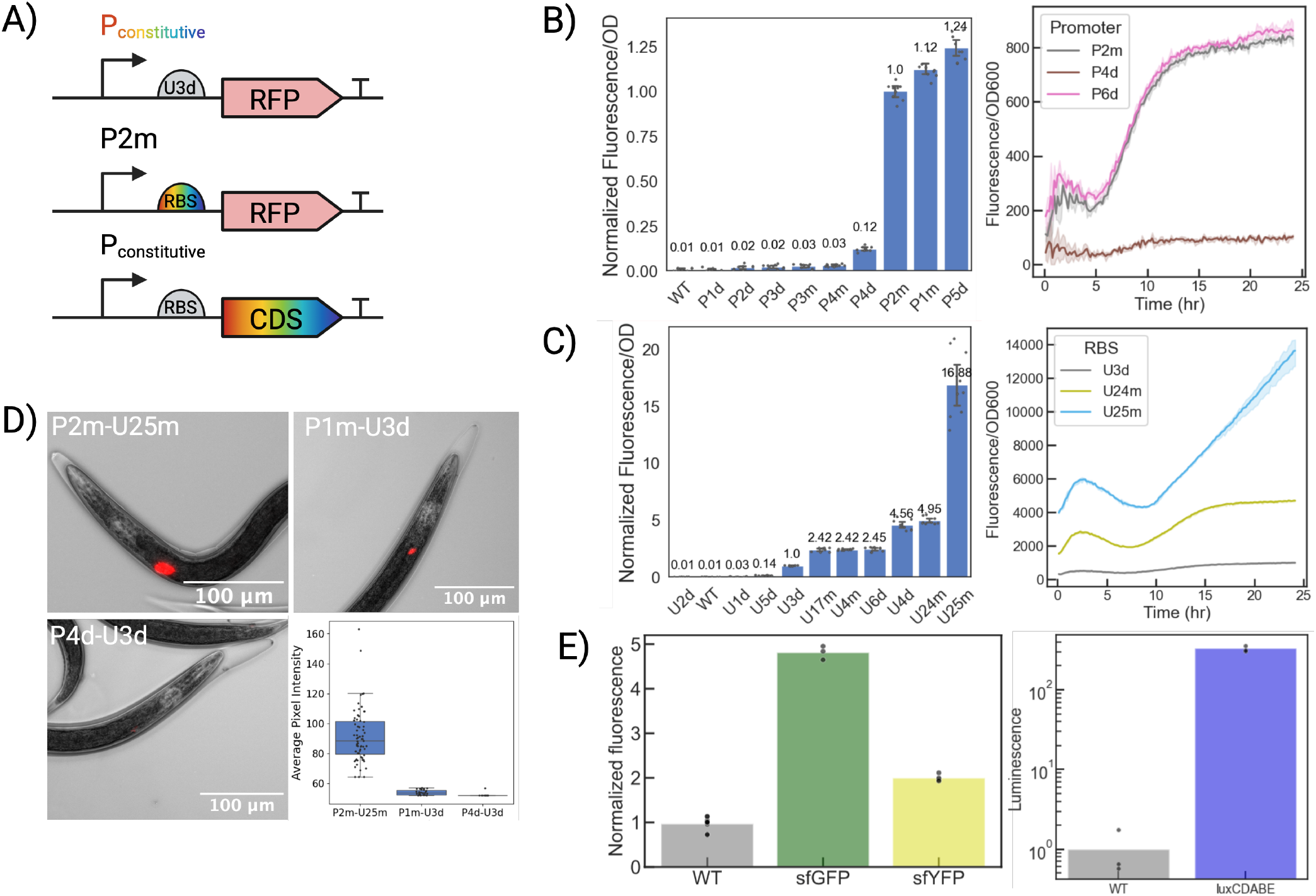
(A) Circuit diagrams for part library. (B) Left: Promoter strength order at the end of 24 hours of growth. Three technical replicates each for three biological replicates are plotted. The fluorescence/OD for all of the strains was normalized by fluorescence/OD of the P2m-U3d strain. The mean normalized fluorescence is plotted with error bars representing 95% confidence intervals. Right: representative time trace for a subset of the strains (all time traces can be found in Figure S3-S5). The solid line represents the mean normalized fluorescence, while lighter shading represents 95% confidence intervals. (C) Left: RBS strength order at the end of 24 hours of growth. Three technical replicates each for three biological replicates are plotted. The fluorescence/OD for all of the strains was normalized by fluorescence/OD of the P2m-U3d strain. The mean normalized fluorescence is plotted with error bars representing 95% confidence intervals. Right: representative time trace for a subset of the strains (all time traces can be found in Figure S6-S8). The solid line represents the mean normalized fluorescence, while lighter shading represents 95% confidence intervals. (D) Upper left and right, lower left: Representative microscopy images of nematodes with intensity values close to the average intensity. Lower right: Average pixel intensity for nematodes colonized by three different bacterial strains. Each dot represents one nematode colonized by bacteria at a detectable level. (E) Left: Normalized fluorescence for strains expressing sfGFP, and sfYFP. Right: Normalized luminescence for the strain expressing the *luxCDABE* operon. Triplicates from one biological replicate are plotted.

For both promoters and RBS we see an increase in fluorescence early in the time trace (Figure 1B-C). This is caused by fluorescent protein already being present in the cells from the overnight growth, which then gets normalized by the low OD from diluting the cells. When growing cells overnight carrying an inducible TurboRFP construct, this increase is not seen (Figure 2C), since there is no TurboRFP present when the inducer is not added.

**Figure 2.**
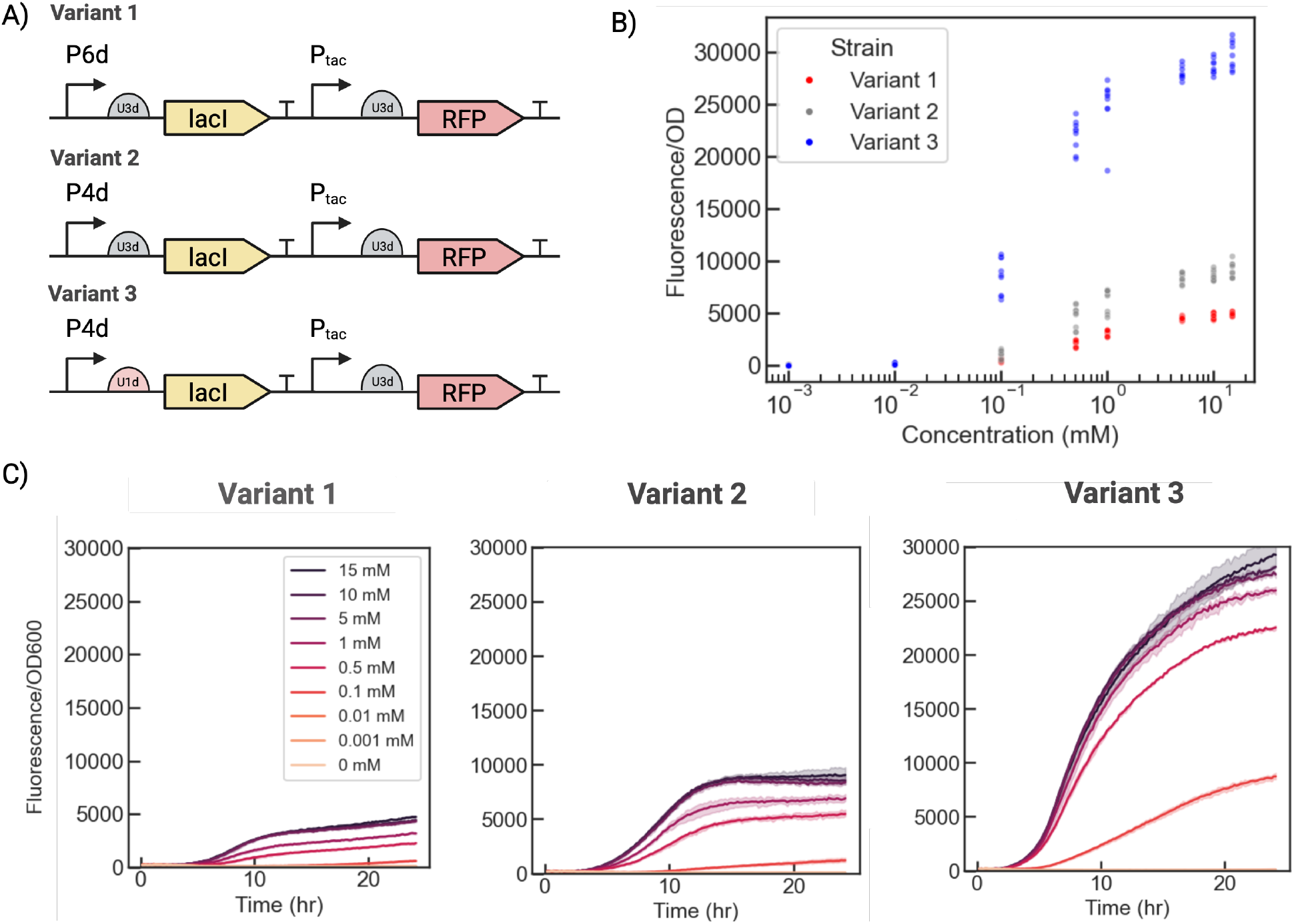
(A) Diagrams for three variants of IPTG inducible constructs. (B) IPTG induction curve for all three strains. Three technical replicates per biological replicate (*n* = 3) are plotted. (C) Time traces for one of the biological replicates for all three strains (all time traces can be found in Figure S10-S12). The solid line represents the mean normalized fluorescence, while lighter shading represents 95% confidence intervals.

In addition to promoter and RBS sequences, we also tested the ability of various coding sequences to express within *X. griffiniae*. For this we tested the expression of sfGFP, sfYFP, and the *luxCDABE* operon. These express well in *Xenorhabdus* (Figure 1E) and offer an alternative readout to the fluorescent proteins that were already known to work well. We also tested mScarlet3, but expression was much weaker than for the previously used TurboRFP (Figure S1).

After characterizing the parts in liquid culture, we chose three strains with different levels of TurboRFP expression to colonize nematodes and measure the fluorescence *in vivo* (Figure 1D). We can see the difference between a strong and weak promoter *in vivo* by imaging the whole nematode and comparing the fluorescence signal (Figure 1D). As predicted, P2m-U25m is stronger than P1m-U3d, but less so than in the liquid culture experiment, probably because the fluorescence signal in the microscopy experiment is saturated for P2m-U25m, or because *in vivo* expression is weaker than *in vitro* expression.

### Design of an IPTG inducible construct

In order to determine whether the characterization data for the constitutive promoters and RBS sequences lead to more predictable performance of a larger genetic circuit, we built a simple IPTG inducible construct (Figure 2A) where we varied the promoters driving expression of *lacI* and the RBS upstream of *lacI* or TurboRFP. All three variants show low leak, meaning that there is enough *lacI* to suppress TurboRFP experssion at the non-induced condition. For Variant 2 and 3, a weaker promoter, P4d (J23106), was chosen to drive expression of *lacI*, leading to increased TurboRFP expression (Figure 2B). The highest TurboRFP expression was achieved for Variant 3 that additionally had a weak RBS, U1d, upstream of *lacI*, resulting in even less *lacI* expression. To quantify the differences between the variants, we fit the data to a Hill equation (estimated parameters for each replicate can be found in Supplemental Information I). The average maximum fluorescence values and *K* values are reported in Table 1.

**Table 1:** The average of the estimated parameter values for maximum fluorescence and *K* values across three biological replicates each having three technical replicates.

| Strain | Max Fluorescence/OD600 | $K$ value |
| --- | --- | --- |
| Variant 1 | 4907 | 0.66 mM |
| Variant 2 | 9035 | 0.54 mM |
| Variant 3 | 29277 | 0.22 mM |

## Materials and Methods

### Cloning

The part library DNA constructs were constructed using the Golden Gate-Gibson (3G) assembly method [13]. Using Golden Gate assembly, the constructs were assembled and and PCR amplified as linear DNA constructs. The linear constructs were inserted into the EML002 backbone (Addgene) a modified plasmid made from of HGB1262 plasmid [8], by Gibson assembly. The Gibson product was then transformed into pir+ competent *E. coli* cells. The transformants were plated on selective (carbenicillin, 100 *µ*g/ml or kanamycin, 50 *µ*g/ml) plates. The chosen colonies were sequence verified by colony PCR. Plasmids were isolated using a Qiagen Miniprep kit. The purified plasmid was then transformed into a diaminopimelic acid (DAP) deficient pir+ *E. coli* strain. The transformants were plated on selective plates with added DAP (0.3 mM).

### Conjugation

The protocol used for conjugation was modified from Alani et al. [17]. Conjugation included three strains: donor *E. coli* strain with the designed DNA construct, helper *E. coli* strain, and receiver *X. griffiniae* HGB2511 strain. The helper *E. coli* strain carries the pUX-BF13 plasmid [18], with *tns* genes. Then, the constructs from the *E. coli* donor strain were integrated as a single copy into the *X. griffiniae* at the Tn7 site downstream of the *glmS* gene. First, the colonies were picked from the DAP deficient donor and helper *X. griffiniae* strains and the *X. griffiniae* strain and were grown in liquid overnights with appropriate antibiotics and DAP added. Wildtype *X. griffiniae* was grown without antibiotics. The next day, the optical density of the liquid cultures were measured so the correct volumes could be pipetted to have an OD of 3. Then, the liquid cultures were washed twice to get rid of remaining antibiotics. Next, the three strains were mixed together and plated onto a filter on LB+DAP agar plates and incubated overnight at 30^*°*^C. The next day, the bacteria were resuspended by vigorously vortexing the filter in a falcon tube filled with 5 ml LB. The conjugation mixture was then plated on kanamycin plates. After a day of growth, *X. griffiniae* colonies were picked and grown in 30 *µ*L LB overnight. The following day, 4 *µ*l of overnight culture was diluted into 46 *µ*l of nuclease free water and boiled for 10 minutes at 100^*°*^C. The boiled culture was then incubated at 4^*°*^C for 10 minutes and centrifuged using a table top centrifuge for two minutes. Then, the colonies were sequence verified using colony PCR (Supplemental Information A).

### Plate reader operations

Starting from an single colony, strains were grown overnight at 30^*°*^C in LB. The overnight cultures were diluted with M9 to OD of 0.05. For the IPTG induction experiments, IPTG was added to the media ranging from 0-15 mM (higher concentrations of IPTG caused a growth defect, see Figure S9). The fluorescence and optical density (600 nm) was measured using a BioTek Synergy H1F Multi-Mode Microplate Reader for 24 hours. TurboRFP fluorescence was measured at an excitation of 550 nm and an emission of 580 nm with a gain of 61 and 100 for the promoter screen and gain of 61 and 80 for the RBS screen and inducible constructs. The cultures were grown in 30^*°*^C and shaken linearly continuously at a frequency of 567 cpm.

### Parameter estimates for IPTG induction curves

The data from the induction curves at the end point after 24 h of growth was fitted, using the curve_fit function in the Python scipy.optimize module [19], to a Hill equation:

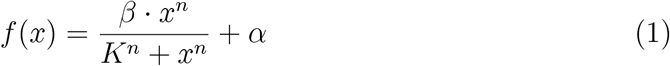

where *β* is the maximum normalized fluorescence, *K* is the inducer concentration at half of the maximum fluorescence, *x* is the inducer concentration, *n* is the Hill coefficient, and *α* is the floor term (accounting for measurement background/leak).

### Preparation of liver-kidney plates

This recipe is a modification of protocol described by Sicard et al. [9] provided by the Cao lab at Carnegie Institute for Science.

For 500 ml agar:

- 50 g beef kidney
- 50 g beef liver
- 2.5 g NaCl
- 7.5 agar
- Water to 500 ml

Use a blender to grind the liver and kidney. Add 250 ml of water and the remaining ingredients to the blender. Blend until almost smooth, but still containing some chunks. Pour into a 1 L flask. Use the remaining water to rinse the blender and pour into the flask to a total final volume of 500 ml. Autoclave the agar for 45 minutes and cool in 55^*°*^C for 30 minutes. Add kanamycin (50 *µ*g/ml). When pouring the plates, swirl the flask in between plates to prevent the chunks from sinking to the bottom.

### Nematode colonization

The nematode colonization protocol was modified from St. Thomas et al. [8]. 700 *µ*L of the bacterial overnight *X. griffiniae* culture was added to a liver kidney (LK) agar plate. Then, 1 mL of axenic *S. hermaphroditum* (PS9179) nematodes was added to each 1.5 mL Eppendorf tube. Axenic nematodes were rendered as described by Cao et al. [5], with the modification of using beef instead of pork liver and kidney. The nematodes were centrifuged for 30 s at 6.0 rcf. Then, 900 *µ*L of the supernatant was discarded and 900 *µ*L of 1% bleach solution was added. The tube was inverted for 1:30 min to kill bacteria on the nematodes’ surface. The nematodes were then washed with water three times to remove residual bleach. Using a dissecting microscope, the number of nematodes in 2 *µ*l droplets was counted to determine the concentration of nematodes in the solution. The appropriate volume of nematodes was then added to each LK plate such that there were approximately 200 nematodes per plate. After one week, the nematodes were trapped in water by placing the LK plate in a larger petri dish where the bottom was covered in autoclaved water. After another week, nematodes were collected from the water traps.

### *In vivo* imaging of nematodes

Before imaging, the nematodes were paralyzed with levamisole (200 *µ*M). The colonized nematodes were imaged using a Nikon Ti2 fluorescence microscope using phase-contrast and the RFP channel at 10x magnification. The RFP channel images went through the same enhancing, denoising, and normalization to adjust the pixel to a range of 1 to 255 (Supplemental Information G). Clusters of at least 150 pixels and a threshold of intensity 40 were identified as colonized regions and cross-referenced with the corresponding phase-contrast nematode images. The average pixel intensities for each colony was calculated by averaging the selected pixels in the determined regions. The percent of detected events was calculated by dividing the number of detected events by the total number of nematodes imaged for each strain (Figure S16).

## Author contributions

EML conceived the idea; EML and OYW designed and performed the experiments, analyzed and visualized the data and wrote the manuscript. EML, OYW and RMM edited the manuscript.

## Acknowledgements

We thank Dr. David Garcia for help with writing the code for the image analysis. We thank Dr. Mengyi Cao for discussions on nematode-bacteria symbiosis, genetics of *Xenorhabdus* bacteria, and the biology of *Steinernema* nematodes, and the Cao group at Carnegie Institution for Science for providing strains, laboratory support, and training on nematode colonization assays. We thank Dr. Mengyi Cao and Carly Myers for maintaining, breeding, and managing the nematodes used in this research. We thank Prof. Heidi Goodrich-Blair for providing the HGB1262 plasmid used in this study. We thank Prof. Dianne Newman for use of the Newman lab fluorescence microscope. All figures were created using BioRender.com. This work was supported by Caltech’s Resnick Sustainability Institute, DaRin Butz Foundation, and Caltech Summer Undergraduate Research Fellowships (SURF) program.

## Supporting Information

All DNA sequences for the designed constructs were sent to Addgene (#85865, plasmid 238378-238408). Supporting Information contains information about primer sequences and engineered strains used in this study, and the assembly processes of these constructs. Supplemental figures for expressed proteins in *X. griffiniae*, time traces for the RBS, promoter, and IPTG screens, and more information regarding the image analysis pipeline is included.

