## Supplemental_information for "A DNA part library for reliable engineering of the emerging model nematode symbiotic bacterium *Xenorhabdus griffiniae* HGB2511"

#### A. Primer sequences for colony PCR and amplification of the EML002 vector

| Primer ID | Sequence | Comment |
|---|---|---|
| oEL245 | GACGTCTAAGAAACCATTATTATCATGAC | Forward primer paired with oEL244 for checking insertion at the Tn7 site in Xenorhabdus, or paired with UNSX_rev for checking donor plasmid |
| oEL244 | TGTCTTACCATGTTGCCTTGA | Reverse primer paired with oEL245 for checking insertion at Tn7 site in Xenorhabdus |
| UNSX_rev | GGTGGAAGGGCTCGGAGTTGTGGTAATCTATGTATCCTGG | UNSX reverse primer |
| UNS1_fwd | CATTACTCGCATCCATTCTCAGGCTGTCTCGTCTCGTCTC | UNS1 forward primer |
| UNSX_fwd | CCAGGATACATAGATTACCACAACTCCGAGCCCTTCCACC | UNSX forward primer |
| UNS1_rev | GAGACGAGACGAGACAGCCTGAGAATGGATGCGAGTAATG | UNS1 reverse primer |
| oEL172 | CTGCTTACATAAACAGTAATACAAGGGGTG | Pair with UNS1_rev for making part 1 of linear backbone of EML002 |
| oEL173 | CACCCCTTGTATTACTGTTTATGTAAGCAG | Pair with UNSX_fwd for making part 2 of linear backbone of EML002 |

#### B. Strain table

| Strain Name | Part | Reference |
|---|---|---|
| HGB1262 | Tn7 vector modified to make backbone for Gibson assembly | St Thomas et al. [8] |
| GYC53 (pUX-B13) | Tns helper plasmid in *E. coli* MFDpir strain background. | Strain provided by McFall-Ngai lab at Carnegie. Plasmid: Bao et al. [18] |
| xOYW1 | UNS1-P1d-U3d-TurboRFP-t0-UNSX | CIDAR MoClo Parts Library [15] |
| xOYW2 | UNS1-P1m-U3d-TurboRFP-t0-UNSX | CIDAR MoClo Parts Library |
| xOYW3 | UNS1-P2d-U3d-TurboRFP-t0-UNSX | CIDAR MoClo Parts Library |

| | | |
|---|---|---|
| xOYW4 | UNS1-P2m-U3d-TurboRFP-t0-UNSX | CIDAR MoClo Parts Library |
| xOYW5 | UNS1-P3d-U3d-TurboRFP-t0-UNSX | CIDAR MoClo Parts Library |
| xOYW6 | UNS1-P3m-U3d-TurboRFP-t0-UNSX | CIDAR MoClo Parts Library |
| xOYW7 | UNS1-P4d-U3d-TurboRFP-t0-UNSX | CIDAR MoClo Parts Library |
| xOYW8 | UNS1-P4m-U3d-TurboRFP-t0-UNSX | CIDAR MoClo Parts Library |
| xOYW9 | UNS1-P5d-U3d-TurboRFP-t0-UNSX | CIDAR MoClo Parts Library |
| xOYW10 | UNS1-P6d-U3d-TurboRFP-t0-UNSX | CIDAR MoClo Parts Library |
| xOYW12 | UNS1-P2m-U1d-TurboRFP-t0-UNSX | CIDAR MoClo Parts Library |
| xOYW13 | UNS1-P2m-U2d-TurboRFP-t0-UNSX | CIDAR MoClo Parts Library |
| xOYW14 | UNS1-P2m-U4d-TurboRFP-t0-UNSX | CIDAR MoClo Parts Library |
| xOYW15 | UNS1-P2m-U5d-TurboRFP-t0-UNSX | CIDAR MoClo Parts Library |
| xOYW16 | UNS1-P2m-U4m-TurboRFP-t0-UNSX | CIDAR MoClo Parts Library |
| xOYW17 | UNS1-P2m-U6d-TurboRFP-t0-UNSX | CIDAR MoClo Parts Library |
| xOYW18 | UNS1-P2m-U17m-TurboRFP-t0-UNSX | CIDAR MoClo Parts Library |
| xOYW19 | UNS1-P2m-U24m-TurboRFP-t0-UNSX | CIDAR MoClo Parts Library |
| xOYW20 | UNS1-P2m-U25m-TurboRFP-t0-UNSX | CIDAR MoClo Parts Library |
| xOYW21 (IPTG Variant 1) | UNS1-P6d-U3d-LacI-t12m-UNS3-Ptac-U3d-TurboRFP-t0-UNSX | CIDAR MoClo Parts Library |
| xOYW22 (IPTG Variant 2) | UNS1-P4d-U3d-LacI-t12m-UNS3-Ptac-U3d-TurboRFP-t0-UNSX | CIDAR MoClo Parts Library |
| xOYW23 (IPTG Variant 3) | UNS1-P4d-U1d-LacI-t12m-UNS3-Ptac-U3d-TurboRFP-t0-UNSX | CIDAR MoClo Parts Library |
| xOYW023 (Xg lum) | UNS1-P2m-U3d-LuxCDABE-t0-UNSX | CIDAR MoClo Parts Library |
| xEML021 | UNS1-P6d-U3d-sfGFP-t0-UNSX | CIDAR MoClo Parts Library |
| xEML022 | UNS1-P6d-U3d-mScarlet3-t0-UNSX | CIDAR MoClo Parts Library |
| xEML046 | UNS1-P6d-U3d-sfYFP-t0-UNSX | CIDAR MoClo Parts Library |

#### C. More information about the construction of the EML002 vector

We removed the chloramphenicol and streptomycin resistance cassettes to allow for more available antibiotic options in deletion strain backgrounds as well as increasing the transformation efficiency. Since two fluorescent proteins, GFPmut3 and TurboRFP, are known to express well in *X. griffiniae* (St Thomas et al., 2024), we made a part plasmid for TurboRFP, where a BsaI cut site in the middle of the coding region was removed. The GFPmut3 part plasmid is already a part of the CIDAR MoClo kit.

#### D. More information about Gibson assembly

We had more success when splitting the plasmid backbone into two pieces and doing a three piece Gibson assembly. Therefore we are using two primer pairs to make backbone pieces from the EML002 plasmid (Supplemental Information A).

#### E. mScarlet and TurboRFP Fluorescence in *X. griffiniae*

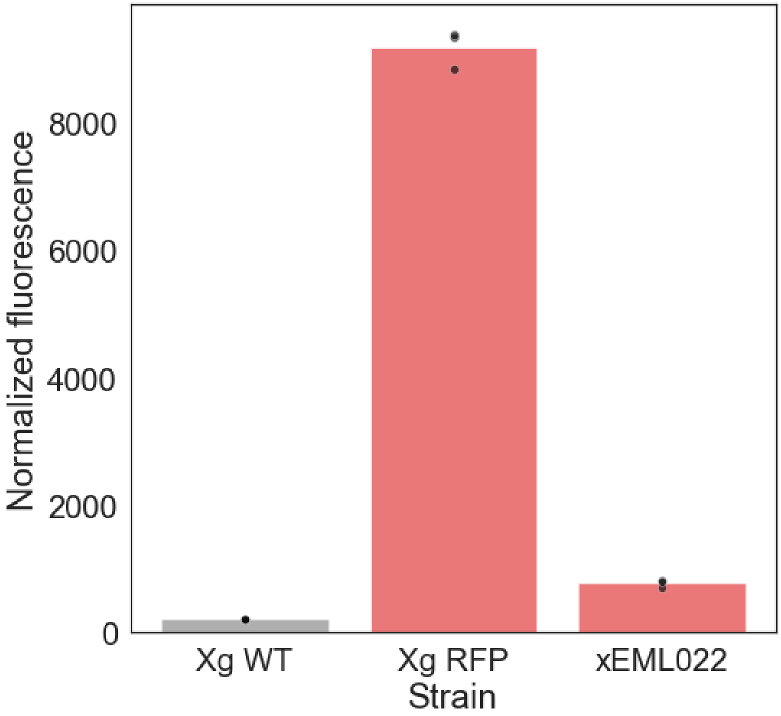


**Figure S1.** Normalized fluorescence for fluorescent proteins expressed in *X. griffiniae*. Three technical replicates of each strain were plotted. TurboRFP had a significantly higher fluorescent signal than mScarlet3.

### F. Time traces from promoter and RBS screen

All of the following figures contain one biological trial with 3 technical replicates.

##### F1. OD curves for Promoter (left) and RBS (right) TurboRFP time curves in Figure 1

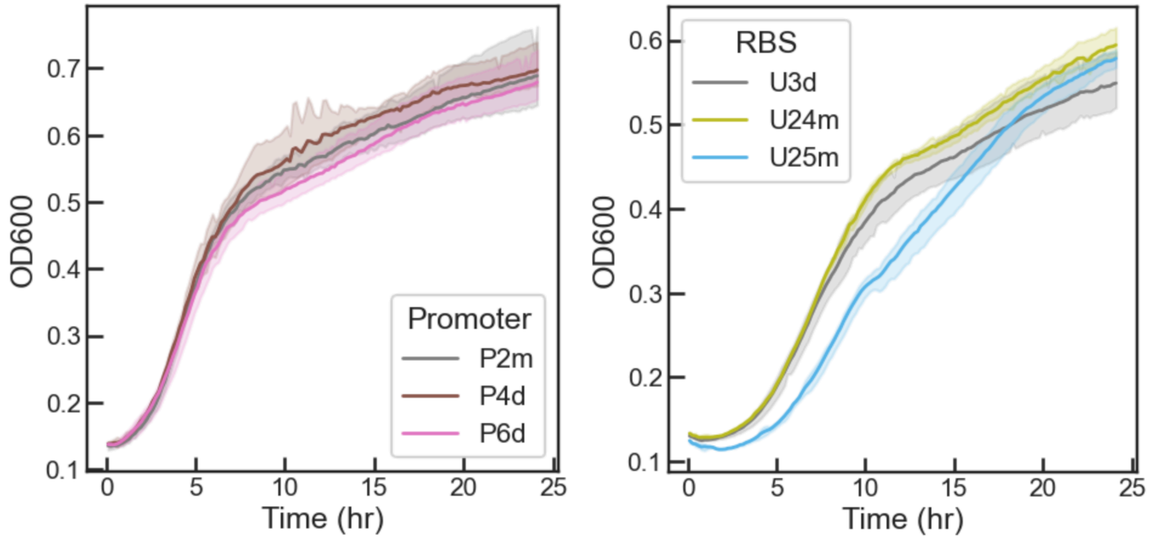


**Figure S2.** OD600 curves for Promoter (left) and RBS (right) strains included in Figure 1B and Figure 1C time curves. The solid line represents the mean normalized fluorescence for each strain, while lighter shading represents 95% confidence intervals.

##### F2. TurboRFP (left) and OD (right) curves for all Constitutive Promoter Strains

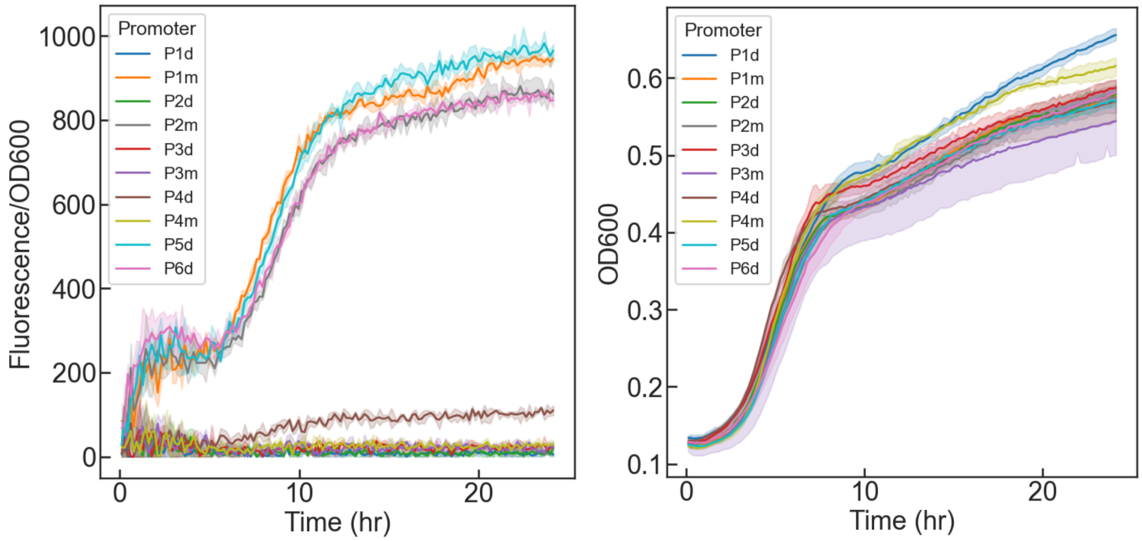


**Figure S3.** Normalized TurboRFP time trace (left) and OD600 curves (right) for all promoter strains in Trial 1. The solid line represents the mean normalized fluorescence for each strain, while lighter shading represents 95% confidence intervals. Because the constitutive strains were grown overnight in LB, there is already TurboRFP present at high levels in the cells, and they have an initial fluorescence when diluted at time 0 hrs. As the bacterial cells grow, the fluorescence does not increase at the same rate as the OD, leading to the initial bump between 0 to 5 hours in the normalized TurboRFP time trace.

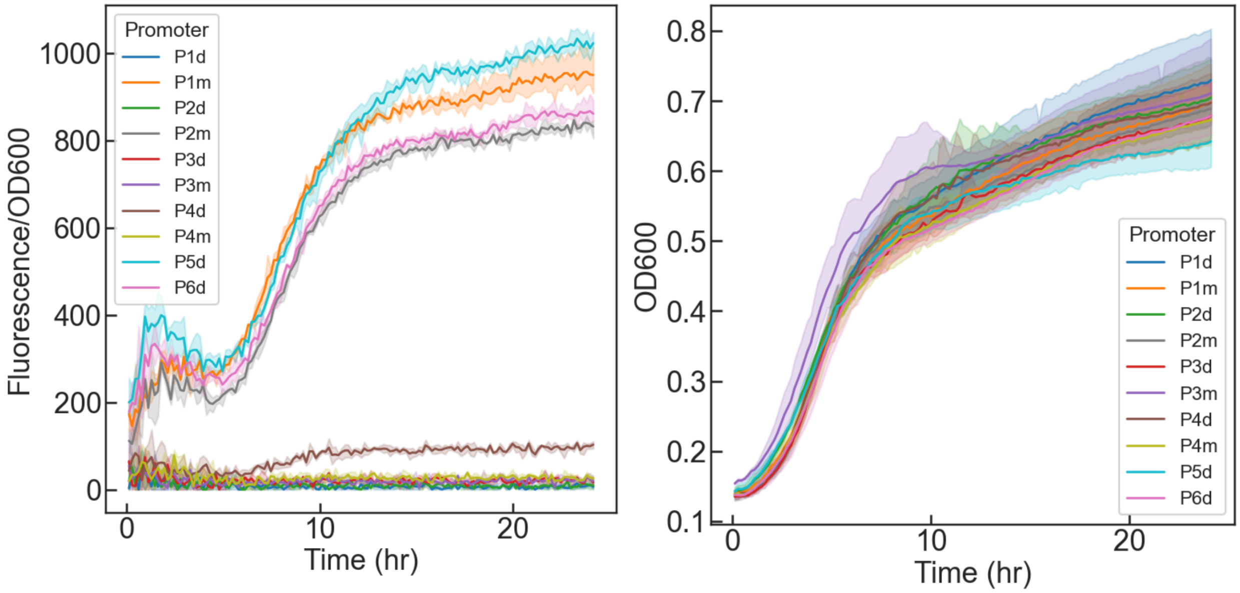


**Figure S4.** Normalized TurboRFP time trace (left) and OD600 curves (right) for all promoter strains in Trial 2. The solid line represents the mean normalized fluorescence for each strain, while lighter shading represents 95% confidence intervals. The initial bump between 0 to 5 hours in the normalized TurboRFP time trace is observed in all trials for promoter strains, described in Figure S3.

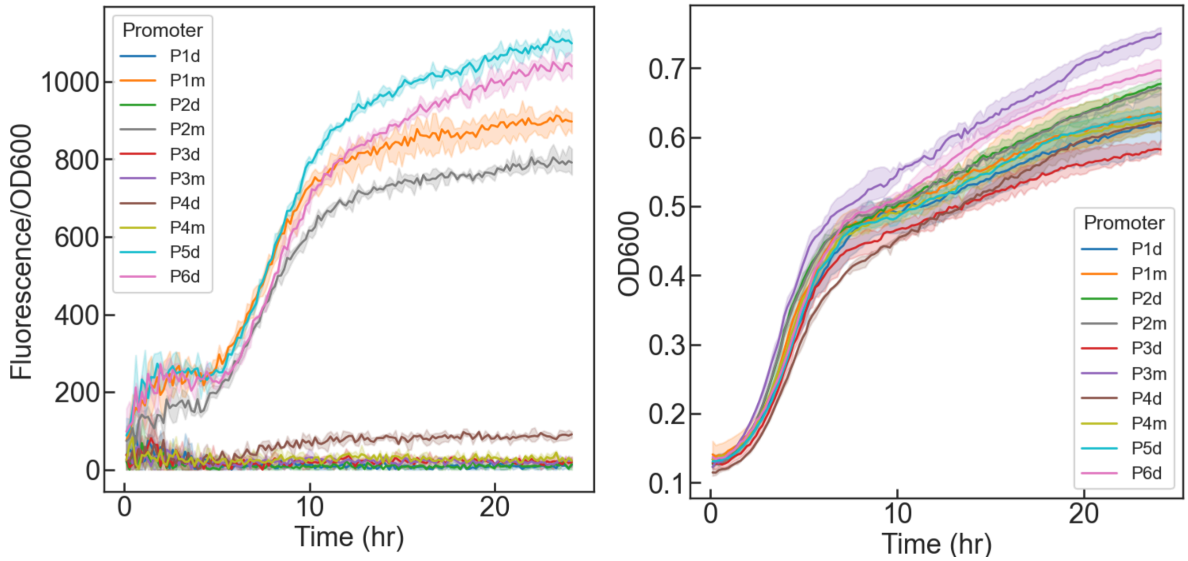


**Figure S5.** Normalized TurboRFP time trace (left) and OD600 curves (right) for all promoter strains in Trial 3. The solid line represents the mean normalized fluorescence for each strain, while lighter shading represents 95% confidence intervals. The initial bump between 0 to 5 hours in the normalized TurboRFP time trace is observed in all trials for promoter strains, described in Figure S3.

**F3. TurboRFP (left) and OD (right) curves for all RBS Strains**

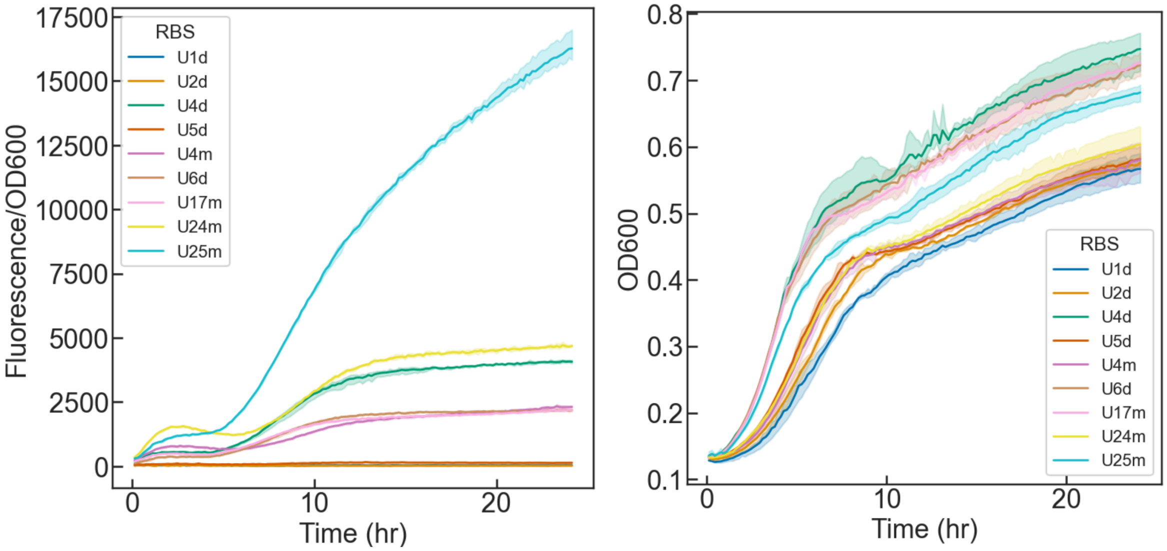


**Figure S6.** Normalized TurboRFP time trace (left) and OD600 curves (right) for all RBS strains in Trial 1. The solid line represents the mean normalized fluorescence for each strain, while lighter shading represents 95% confidence intervals. The initial bump between 0 to 5 hours in the normalized TurboRFP time trace is observed in both promoter (Figure S3) and RBS strains. Because the RBS strains also have constitutive promoters, the cells grown overnight have an initial fluorescence. During the first few hours of the experiment, the cells are dividing quickly, so the fluorescence does not increase at the same rate of the OD. Therefore, a characteristic bump appears in the normalized fluorescence time curve.

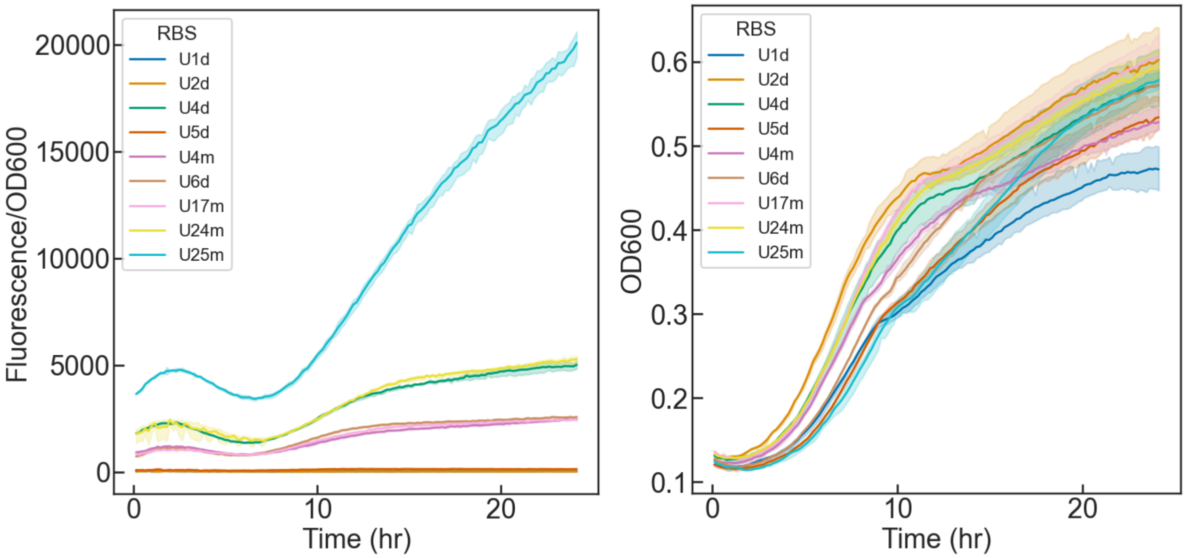


**Figure S7.** Normalized TurboRFP time trace (left) and OD600 curves (right) for all RBS strains in Trial 2. The solid line represents the mean normalized fluorescence for each strain, while lighter shading represents 95% confidence intervals. The initial bump between 0 to 8 hours in the normalized TurboRFP time trace is observed in all trials for RBS strains, described in Figure S6.

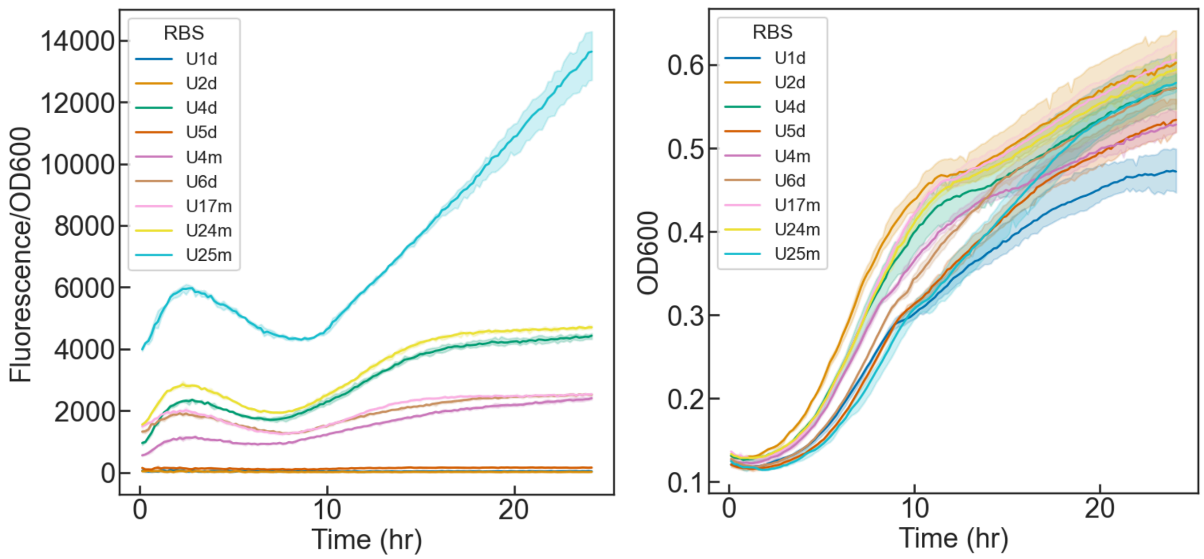


**Figure S8.** Normalized TurboRFP time trace (left) and OD600 curves (right) for all RBS strains in Trial 3. The solid line represents the mean normalized fluorescence for each strain, while lighter shading represents 95% confidence intervals. The initial bump between 0 to 9 hours in the normalized TurboRFP time trace is observed in all trials for RBS strains, described in Figure S6.

#### G. Information about image analysis pipeline

Please refer to the attached supplemental file for image analysis code.

#### H. Colony and nematode counts in microscope images

The table below describes the number of detected colonization events per strain in the microscopy experiment as well as how many nematodes were counted per strain.

| Strain | Colony Counts | Nematode Counts |
|---|---|---|
| P1m-U3d | 26 | 37 |
| P4d-U3d | 11 | 54 |
| P2m-U25m | 32 | 94 |

### I. Time traces inducible constructs

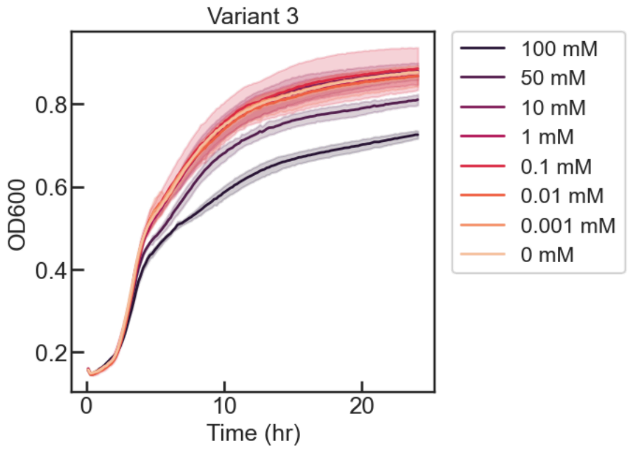


**Figure S9.** IPTG induction curve for finding limit before the inducer concentration becomes toxic, resulting in a growth penalty.

All of the following figures contain one biological trial with 3 technical replicates.

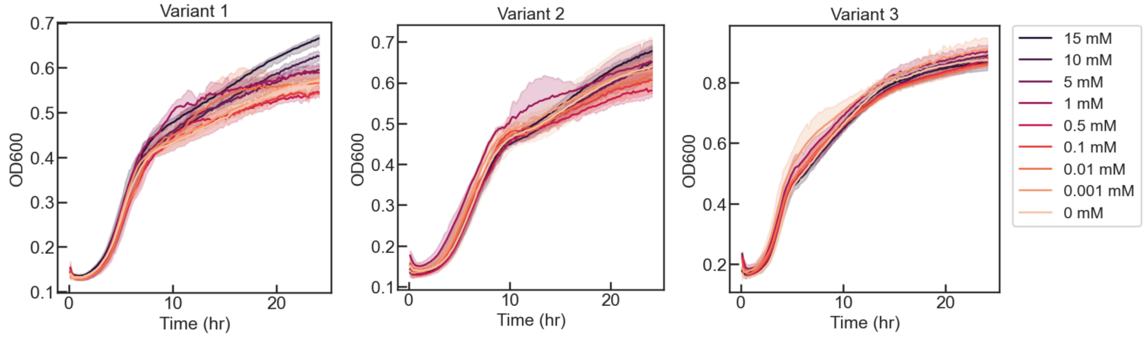


**Figure S10.** OD600 curves for each inducible variant (Left: Variant 1, Center: Variant 2, Right: Variant 3) in Trial 1. Normalized TurboRFP time curves in Figure 2. The solid line represents the mean normalized fluorescence for each IPTG concentration, while lighter shading represents 95% confidence intervals.

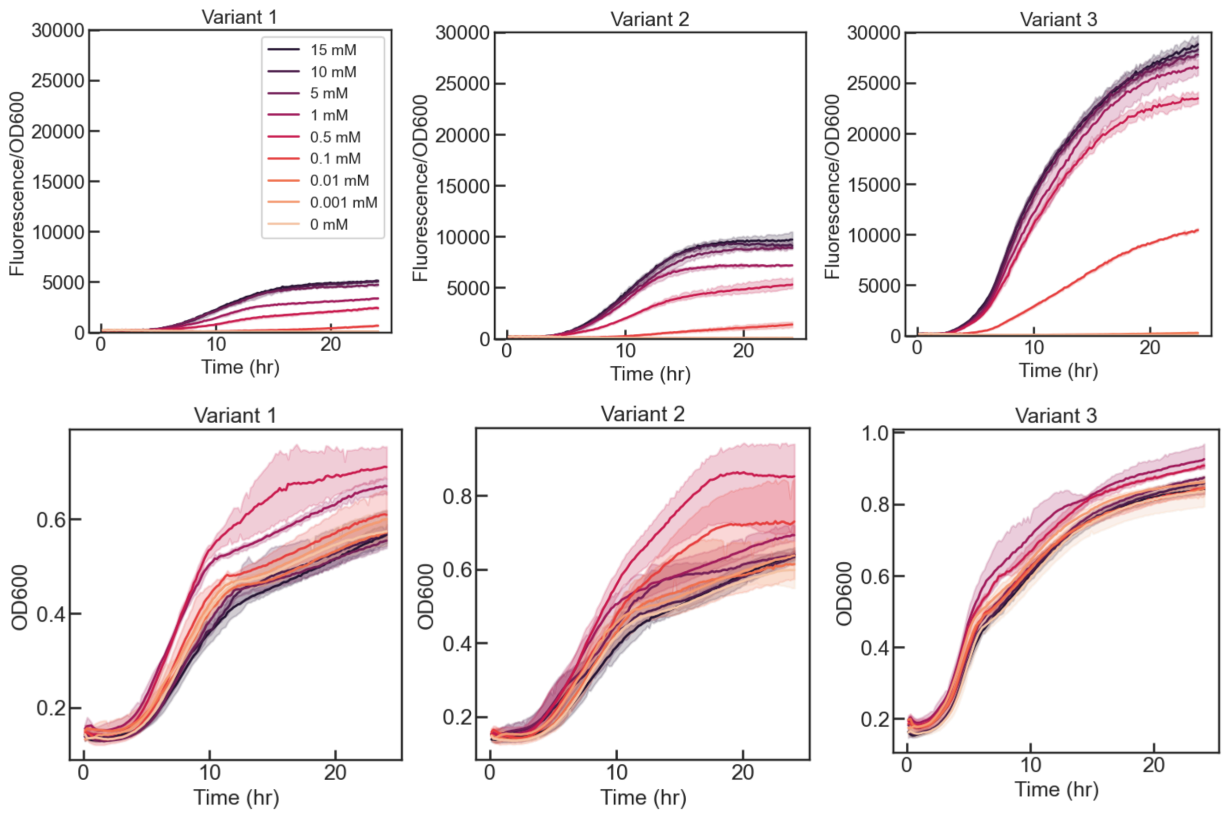


**Figure S11.** Normalized TurboRFP (top) time traces and OD (bottom) curves for each inducible variant (Left: Variant 1, Center: Variant 2, Right: Variant 3) in Trial 2. The solid line represents the mean normalized fluorescence for each IPTG concentration, while lighter shading represents 95% confidence intervals. The initial bump observed in both promoter and RBS strains does not occur here because the variants have an IPTG inducible promoter. When the variants are grown overnight and diluted for the experiment they do not have an initial fluorescence. Therefore, as the cells grow and divide in the first 5 hours, the normalized fluorescence stays at around 0. After 5 hours, the normalized fluorescence curve begins to increase and differentiate based on IPTG concentration.

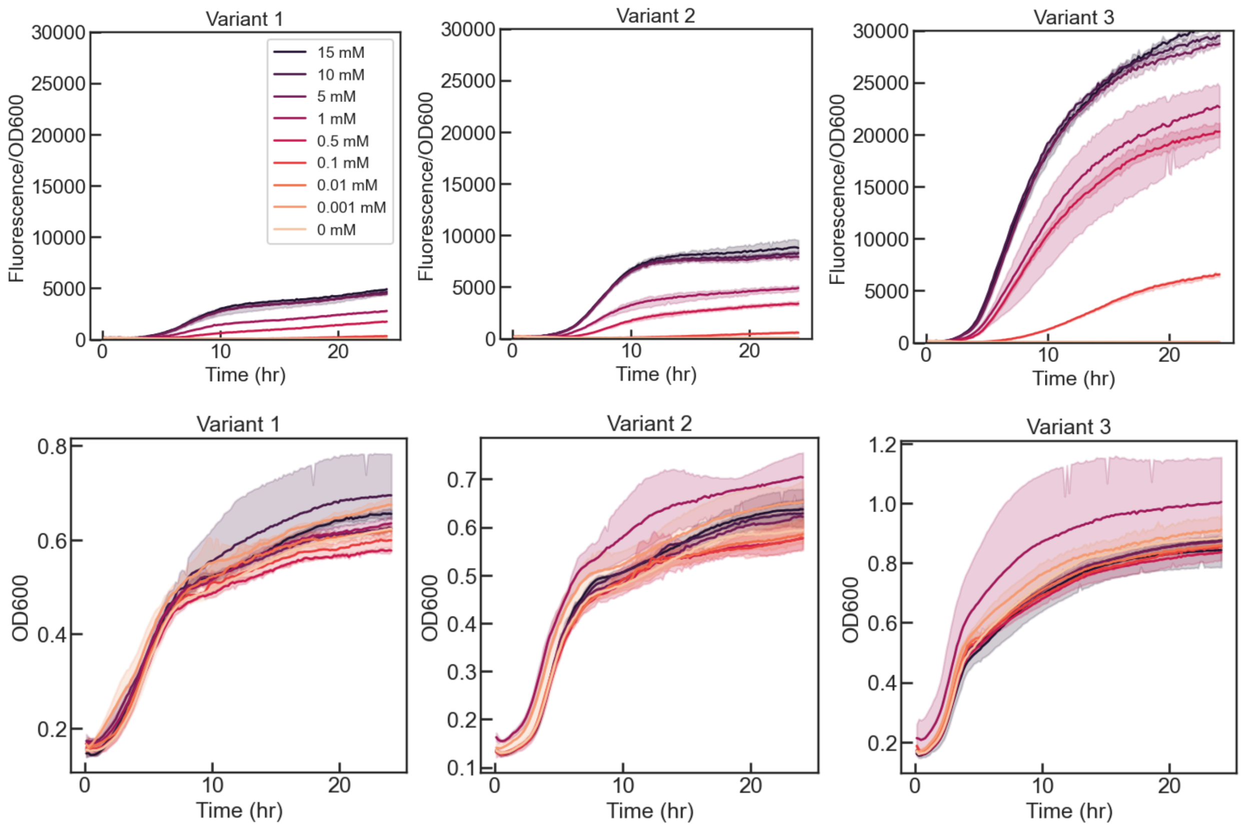


**Figure S12.** Normalized TurboRFP (top) time traces and OD (bottom) curves for each inducible variant (Left: Variant 1, Center: Variant 2, Right: Variant 3) in Trial 3. The solid line represents the mean normalized fluorescence for each IPTG concentration, while lighter shading represents 95% confidence intervals.

**Table S1**: Estimated parameters for each replicate in the IPTG induction experiment.

| Strain | Biological replicate | Technical replicate | β | K | n | α |
|---|---|---|---|---|---|---|
| Variant 1 | 1 | 1 | 4640.7113 | 0.55717831 | 1.212869 | 51.7696489 |
| Variant 1 | 1 | 2 | 4643.10486 | 0.54748303 | 1.1780457 | 53.6020738 |
| Variant 1 | 1 | 3 | 4587.59175 | 0.53147028 | 1.14309386 | 39.4801257 |
| Variant 1 | 2 | 1 | 5067.61907 | 0.55322075 | 1.1664078 | 46.6582014 |
| Variant 1 | 2 | 2 | 5279.92126 | 0.64871055 | 1.10374586 | 47.3813316 |
| Variant 1 | 2 | 3 | 5287.2217 | 0.61012123 | 1.02788448 | 40.5908637 |
| Variant 1 | 3 | 1 | 4951.77718 | 0.8395324 | 1.27376762 | 59.2512756 |
| Variant 1 | 3 | 2 | 4855.84627 | 0.83133782 | 1.2298046 | 45.0235237 |
| Variant 1 | 3 | 3 | 4853.41979 | 0.81894065 | 1.26409289 | 43.7292673 |
| Variant 2 | 1 | 1 | 9274.03927 | 0.35223695 | 1.28471025 | 8.81554742 |
| Variant 2 | 1 | 2 | 8632.04629 | 0.37763571 | 1.33692886 | 11.2483529 |
| Variant 2 | 1 | 3 | 8357.71386 | 0.36650455 | 1.48116975 | 42.5286552 |
| Variant 2 | 2 | 1 | 9949.0652 | 0.38875063 | 1.0725657 | 2.81E-25 |
| Variant 2 | 2 | 2 | 9429.60861 | 0.46205244 | 1.35019782 | 53.3330703 |
| Variant 2 | 2 | 3 | 8942.24918 | 0.40608048 | 1.36967964 | 79.9510742 |
| Variant 2 | 3 | 1 | 9579.60256 | 0.85003153 | 1.05525806 | 5.58E-17 |
| Variant 2 | 3 | 2 | 8518.18832 | 0.77633929 | 1.26200055 | 29.2053513 |
| Variant 2 | 3 | 3 | 8629.01947 | 0.84359092 | 1.14799182 | 18.5159612 |
| Variant 3 | 1 | 1 | 27963.6369 | 0.18205145 | 1.40547392 | 1.40E-17 |
| Variant 3 | 1 | 2 | 28452.205 | 0.18720572 | 1.34531371 | 4.90E-19 |
| Variant 3 | 1 | 3 | 28956.1149 | 0.18534348 | 1.3046906 | 3.99E-15 |
| Variant 3 | 2 | 1 | 27965.2955 | 0.14521097 | 1.37199647 | 3.83E-14 |
| Variant 3 | 2 | 2 | 28756.9058 | 0.15826533 | 1.32460882 | 1.68E-16 |
| Variant 3 | 2 | 3 | 28632.1345 | 0.15374081 | 1.32742288 | 2.62E-12 |
| Variant 3 | 3 | 1 | 32067.3634 | 0.40323184 | 0.84691886 | 3.38E-13 |
| Variant 3 | 3 | 2 | 30161.1979 | 0.28848591 | 1.19660511 | 3.25E-17 |
| Variant 3 | 3 | 3 | 30535.6164 | 0.30498117 | 1.20041107 | 1.01E-17 |

**Table S2:** Mean of estimated parameters in the IPTG induction experiment.

| Strain | B (mean) | B (standard error of mean) | K (mean) | K (standard error of mean) | N (mean) | n (standard error of mean) | α (mean) | A (standard error of mean) |
|---|---|---|---|---|---|---|---|---|
| Variant 1 | 4907.468131913113 | 88.15231528 | 0.6597772237565843 | 0.044193329220167575 | 1.1777457572358998 | 0.026321613929003382 | 47.49847908592574 | 2.130096107043148 |
| Variant 2 | 9034.614750730536 | 182.82379839231166 | 0.5359136129066999 | 0.07289618 | 1.2622780489961711 | 0.047895137481418555 | 27.06644584539051 | 9.015401510039613 |
| Variant 3 | 29276.718927888767 | 457.4558785270299 | 0.22316852041824534 | 0.029558593331813963 | 1.2581601601488093 | 0.0565043 | 3.333214974784113e-13 | 2.8809974287201216e-13 |

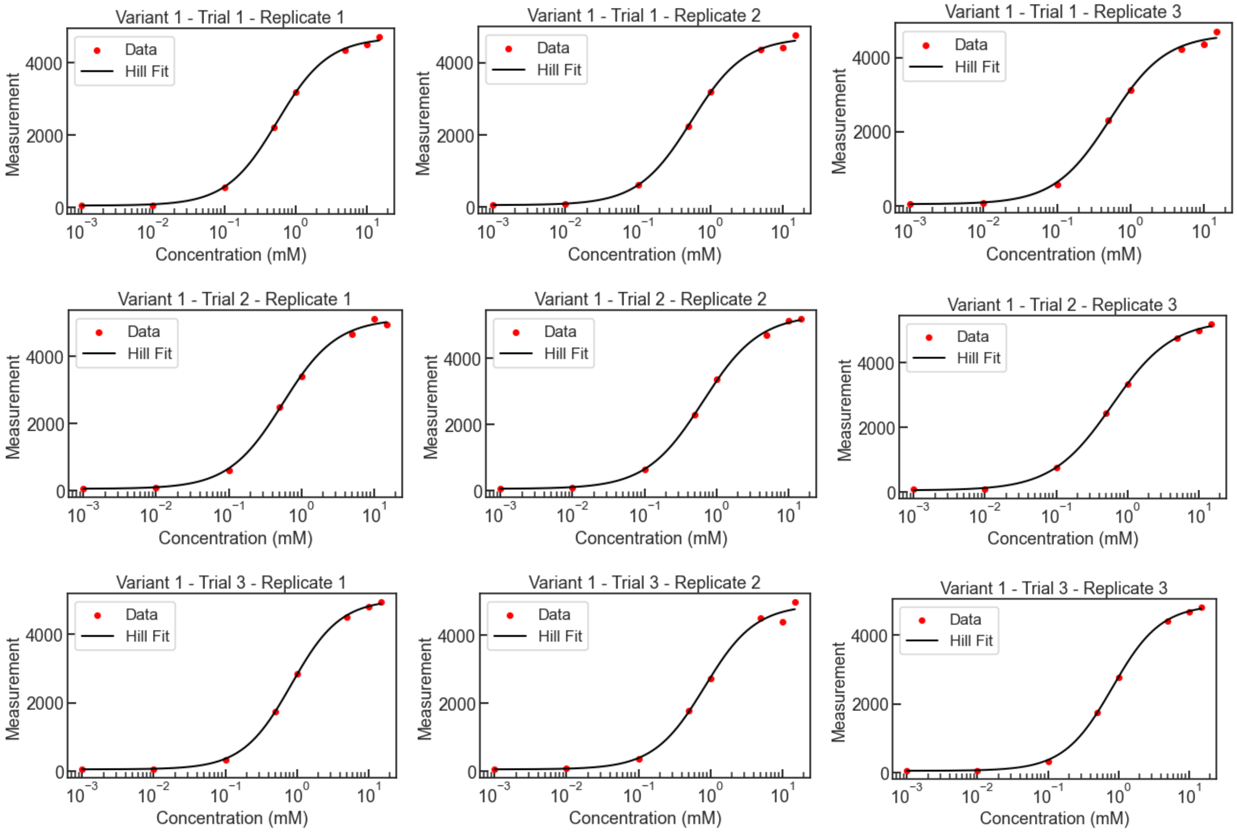


**Figure S13**. Hill Function Curve Fit for Variant 1.

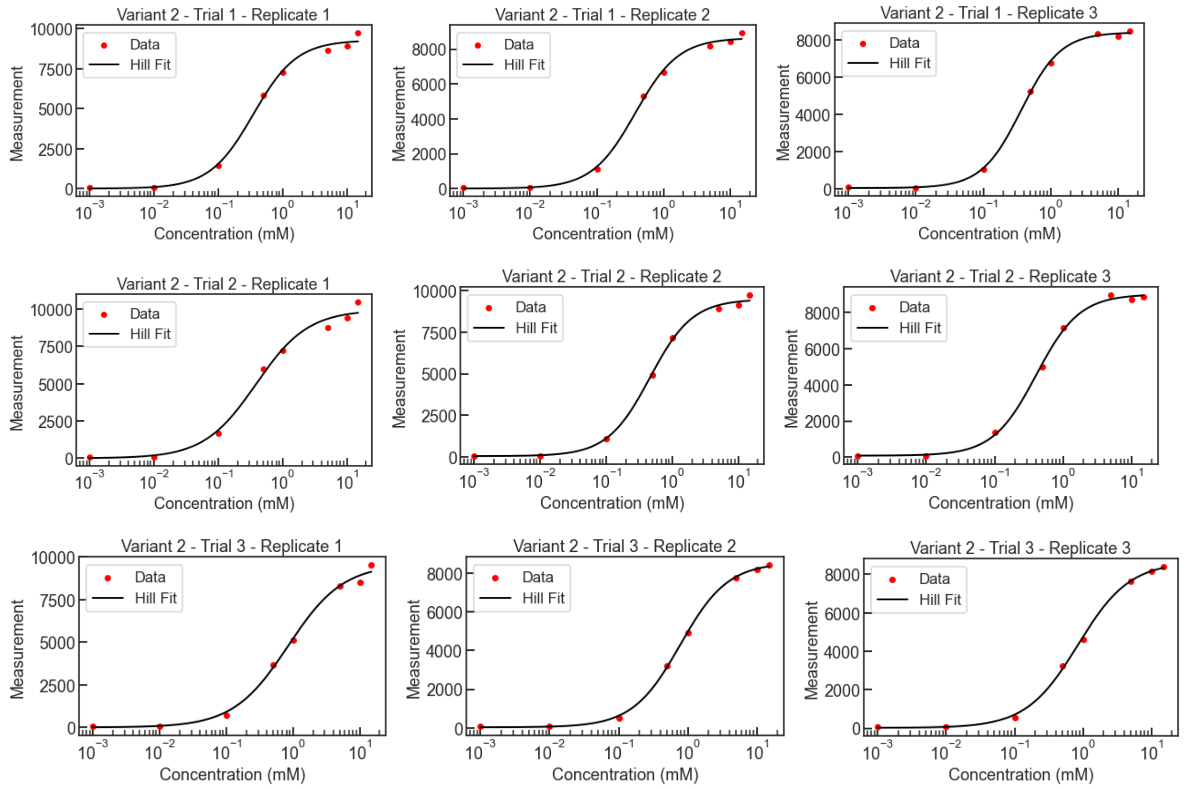


**Figure S14**. Hill Function Curve Fit for Variant 2.

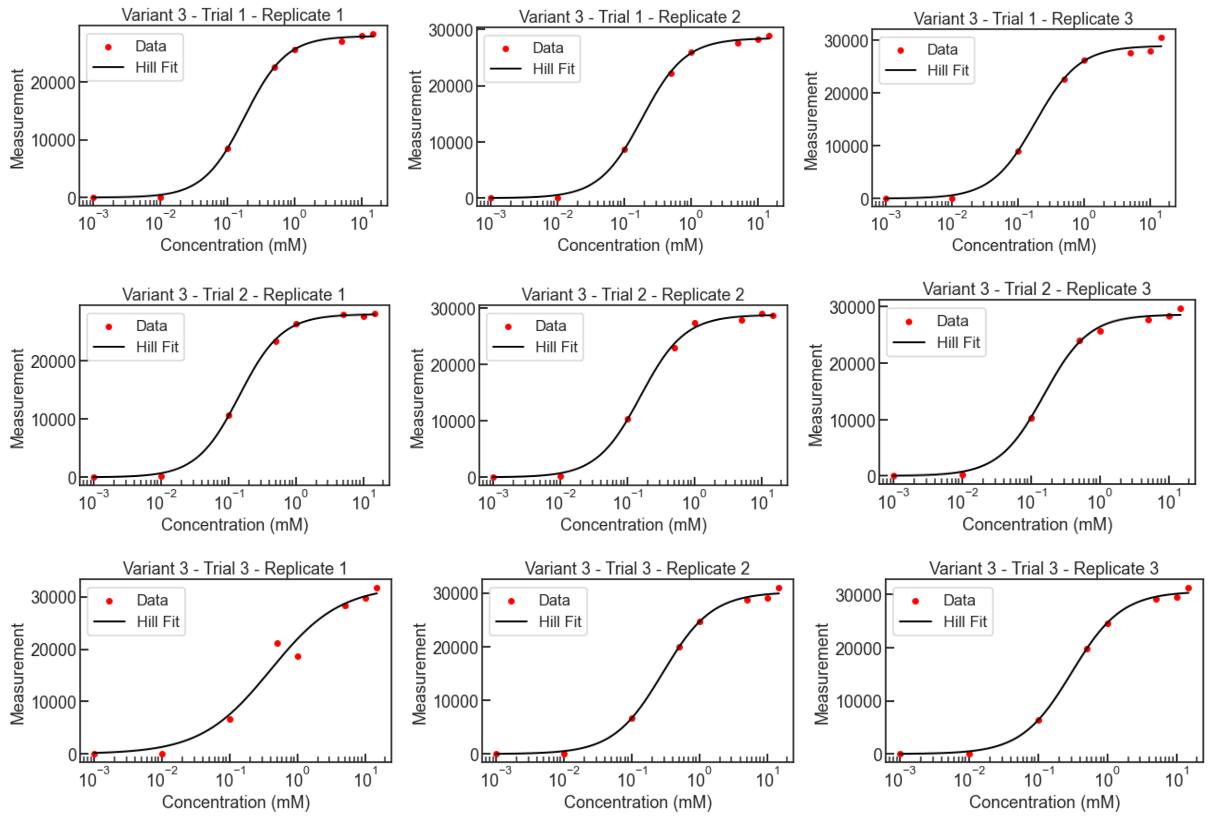


**Figure S15**. Hill Function Curve Fit for Variant 3.

#### J. Additional images from pixel intensity analysis

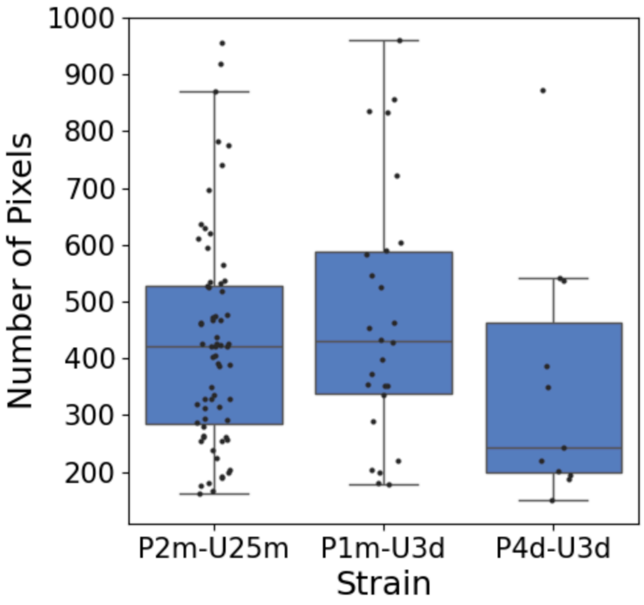


**Figure S16**. Area of bacterial colonization in the nematodes. Each dot represents one detected colonized nematode.
